# Targeting GSK-3β with Peptide Inhibitors: A Rational Computational Strategy for Alzheimer’s Disease Intervention

**DOI:** 10.1101/2024.12.30.630755

**Authors:** Sara Zareei, Saeed Pourmand, Negin Rahimi, Shokoufeh Massahi

## Abstract

Glycogen synthase kinase-3 beta (GSK-3β) is a pivotal serine/threonine kinase implicated in Alzheimer’s disease (AD) pathogenesis, particularly through the hyperphosphorylation of tau protein and increased production of amyloid-beta (Aβ) peptides. This study investigates kappa casein-derived peptides as potential inhibitors of GSK-3β. A peptide library of 42 sequences was generated from kappa casein and docking studies identified IP8 (LRFFVAPFPE) as the top candidate for binging to the previously approved inhibition site of GSK-3β. Molecular dynamics (MD) simulations revealed that IP8’s first three residues contributed unfavorably to binding, promoting the design of mutated peptides (MPs) MP27, MP31, and MP39. Of these, MP31 (HPDFVAPFPE) demonstrated the most stable interaction with GSK-3β, exhibiting the most favorable binding score (-96.6) and interacting with 19 residues of the ATP-binding pocket of the enzyme. Structural analyses confirmed MP31’s superior stability, with minimal RMSD deviations, and stable hydrogen bond formation. The results showed that peptide binding stabilizes GSK-3β by reducing both domain-level dynamics and local side-chain flexibility, leading to a more structurally constrained enzyme. This dual stabilization of the backbone and side chains underscores the critical role of peptide interactions in modulating the conformational landscape of GSK-3β and potentially obstructing substrate access or product release, which are crucial for enzyme activity. These results suggest that kappa casein-derived peptides, particularly MP31, could be promising therapeutic candidates for inhibiting GSK-3β in AD.

## Introduction

The investigation of Glycogen synthase kinase3 beta (GSK-3β), a multifunctional serine/threonine kinase, as a therapeutic target in Alzheimer’s disease (AD) holds significant promise due to its central role in the pathogenesis of the disease. GSK-3β is implicated primarily through its involvement in the hyperphosphorylation of tau protein, a key process that triggers the formation of neurofibrillary tangles (NFTs), which are hallmark features of AD. These NFTs are strongly associated with neuronal dysfunction and synaptic dysfunction, ultimately contributing to neural death and the cognitive decline observed in AD patients (1). Elevated GSK-3β activity is closely linked to abnormal tau aggregation, disrupting neural integrity and accelerating disease progression (2, 3).

Beyond its role in tau hyperphosphorylation, GSK-3β also influences the generation of amyloid-beta (Aβ) peptides, another central feature of AD pathology. By modulating the cleavage of amyloid precursor protein (APP), GSK-3β enhances Aβ production, which contributes to the formation of amyloid plaques in AD brains (4).

Moreover, dysregulated GSK-3β activity is associated with synaptic loss and impaired long-term potentiation (LTP), processes critical for memory and learning. This exacerbates the cognitive decline observed in AD (5), highlighting the importance of GSK-3β in synaptic function and neural communication. Therefore, targeting GSK-3β could help alleviate some of these cognitive impairments.

Additionally, GSK-3β has been shown to negatively regulate insulin signaling, a pathway that plays a critical role in brain function. Impaired insulin signaling and insulin resistance are recognized risk factors for AD, and by modulating GSK-3β activity, it may be possible to restore insulin signaling and potentially mitigate the cognitive decline associated with the disease (6, 7).

In recent years, peptide-based therapeutics have gained significant recognition for addressing various conditions, including cancer (8, 9), diabetes (10), and cardiovascular disease (11). Peptides offer several advantages over small molecules and antibodies, owing to their ease of modification and ability to penetrate tissues and cells. Additionally, their high biocompatibility and low immunogenicity in vivo make them attractive candidates for AD treatment. It is shown that anti-Alzheimer’s peptides have been shown to act through various mechanisms, including the inhibition of Aβ formation and aggregation, mitigation of neuroinflammation, reduction of oxidative stress, and reduction of tau tangles (12–14).

Casein, a milk protein, has demonstrated potential in alleviating cognitive decline in AD mouse models (15). Casein comprises a group of closely related phosphoproteins, each possessing distinct characteristics and functions within the realms of nutrition and food science. The primary types of casein include αS1-Casein, αS2-Casein, β-Casein, κ-Casein, and γ-Casein. Kappa casein (κ-Casein), constituting approximately 10-12% of bovine milk casein, is vital for maintaining the stability of casein micelles by forming a protective layer that prevents micelle aggregation (16).

Despite extensive investigations of therapeutic targets and drug candidates for AD, a critical research gap persists in the limited exploration of kappa casein-derived peptides as potential inhibitors of GSK-3β, which can be considered a promising avenue for therapeutic advancement.

Computational approaches, particularly molecular docking and molecular dynamics simulations (MD) have emerged as indispensable tools in AD research due to their ability to accelerate drug discovery and elucidate its underlying molecular mechanisms. Molecular docking is utilized for virtual screening and drug design by providing a structural insight into the potential affinities for lead optimization (17). Complementing this, MD simulations offer insights into the dynamic stability of protein-ligand complexes over time, capturing conformational changes, and solvent interactions. These simulations help refine binding affinities, reveal structural flexibility, and uncover key residues involved in the regulation or inhibition of key enzymes. Together, these techniques facilitate the rational design of drugs aimed at mitigating the complex, multifactorial nature of AD (18, 19).

In this study, we utilized advanced molecular docking and molecular dynamics simulations to design novel peptide inhibitors derived from food sources, with promising potential for the treatment of Alzheimer’s disease (AD). This study introduces novel peptide modulators for targeting GSK-3β using naturally derived peptides.

## Computational Approaches

### Peptide Library Design

We used kappa casein (Uniprot: P02668) as the parent sequence to generate a peptide library. The fasta sequence of kappa casein was submitted to the Peptide Library Design Tool on the GenScript online server (https://www.genscript.com/peptide_screening_tools.html) to generate 10-length peptides with a 5-residue overlap.

The I-TASSER (Iterative Threading ASSEmbly Refinement) server (https://zhanggroup.org/I-TASSER/) was utilized to predict the 3D conformation of each peptide in the library.

### Peptide-Enzyme Docking

Molecular docking was conducted using the HADDOCK (High Ambiguity Driven Biomolecular DOCKing) server to investigate the inhibitory potential of the peptides on GSK-3β by competing with ATP as a substrate. The 3D structures of the peptides and GSK-3β (PDB ID: 3Q3B, chain A) were input to the server. The lowest-scoring docking clusters were considered the conformation of the peptide-GSK-3β complex.

### Allergenicity and Toxicity Prediction

To ensure ideal anti-Alzheimer’s peptides, it is crucial to eliminate allergenic and toxic peptide candidates. This is because Alzheimer’s disease primarily affects the elderly population, which may be more susceptible to adverse reactions from allergenic or toxic substances. Moreover, Allergenic or toxic peptides can trigger unwanted immune responses, which may reduce the efficacy of the drug and increase the risk of side effects in patients. We used the AllerTOP and ToxinPred servers, which employ k-nearest neighbors’ algorithms and support vector machine (SVM) classifiers to predict allergens and toxic peptides, respectively.

### Molecular Dynamics Simulations

The dynamics of peptide-GSK-3β complexes were evaluated using molecular dynamics simulation. The procedure was similar to our previous study (20) including forcefields, water molecules, equilibration steps, and algorithms. Briefly, the simulations were performed using GROMACS 2019.1. The topology and coordinate files for the protein and peptides were generated using the CHARMM27 force field. Each cubic-shaped system was with extended simple point charge (SPC) water molecules, maintaining a 1 nm buffer from the box edges under periodic boundary conditions. Chloride and sodium ions were added to neutralize the system’s charge.

Energy minimization was performed for 50,000 steps with a 2fs time step, followed by equilibration at 300 K using the Berendsen thermostat and a pressure of 1 bar under NVT and NPT ensembles. The 150 ns production run was conducted using the Particle Mesh Ewald (PME) method to account for long-range electrostatic interactions and the Verlet algorithm for trajectory calculations.

The following analyses were performed: Root-mean-square deviation (RMSD), root mean square fluctuation (RMSF), the radius of gyration (Rg), solvent-accessible surface area (SASA), hydrogen bond analysis, and Molecular mechanics Poisson–Boltzmann surface area (MM-PBSA). Systems underwent a single150ns production step, and the topology coordinates of the peptides were generated using the gmx pdb2gmx built-in tool.

Principal component analysis (PCA) was performed on the protein’s trajectory data, focusing on the last 10 ns (1000 frames) of the simulation. The PCA data was clustered using the K-means algorithm, with the number of clusters determined based on the structural characteristics of the system. For each cluster, the centroid was defined as the average position of all frames within that cluster in the reduced dimensional PCA space (PC1 and PC2). Then, the Euclidian differences were calculated between centroids with all frames and the closest frames to the centroids were determined. Next, RMSD values were computed to evaluate the conformational differences between clusters.

### Mutagenesis

We used the MCSM (Membrane Curvature Server for Molecular Dynamics) server to introduce mutations in the peptide sequence based on the Delta Delta G (ΔΔG) concept. ΔΔG predicts the change in peptide stability caused by a single point mutation and is calculated as the difference in binding free energy between the mutant peptide and wild-type peptides. ΔΔG=ΔG_wild_−ΔG_mutant_

Positive ΔΔG values indicate a stabilizing effect of the mutation, while negative values suggest decreased peptide stability. ΔΔG ≈ 0 is also known as a mutation with no effect on peptide stability.

The MutLib.py script (https://github.com/SaraZareei/Peptide-Design/blob/main/MutLib.py) was used to generate all possible sequence combinations from the favorable mutated residues for each position provided by the MCSM server.

## Results and Discussion

GSK-3β is a promising therapeutic target in mitigating the progression of AD due to its role in key pathological mechanisms such as tau hyperphosphorylation, Aβ production, synaptic dysfunction, and insulin signaling. As a result, the developing GSK-3β-targeted therapies offer a promising avenue in the ongoing search for effective treatments. In light of the proven beneficial effects of milk extract against AD (21), we sought to investigate kappa-casein-derived peptides as potential therapeutic agents.

There is a wide variety of GSK-3β inhibitors related to AD, each differing based on their specific target (22). The most common and diverse category consists of inhibitors that act on a single target/function. This group includes both ATP-competitive and non-ATP competitive inhibitors, at the interface of α-helical and β-sheet domains or inhibit GSK-3β activity through other mechanisms, respectively (23–25).

Another significant category encompasses multifunctional inhibitors (22). These compounds not only inhibit GSK-3β but also offer additional therapeutic benefits by targeting other enzymes or pathways associated with AD. For example, some multifunctional inhibitors also inhibit BACE1 or cholinesterase enzymes (26, 27), which are implicated in amyloid plaque formation. Others act on key AD-related pathways, such as reducing tau phosphorylation (28), alleviating oxidative stress (29), and minimizing neuroinflammation (30).

From a structural view, two distinct domains is seen in GSK-3β: a β-barrel shaped N-terminal domain consisting of the first 134 residues, and the C-terminal composed of α-helices, formed by the remaining residues. GSK-3β has several cavities where various agents can accommodate including ATP binding site, substrate site, Axin/fratide site, as well as four allosteric sites (Figure 1). The design of peptide inhibitors in this study was inspired by the approved inhibitor 4-(4-hydroxy-3-methylphenyl)-6-phenylpyrimidin-2(5H)-one (or 55E). This chemical binds to several key regions of the enzyme, including the ATP binding site (Ala83, Lys85, Asp133, and Val135), the catalytic loop (Asp200), the glycine-rich loop (Gly63, Gly64, Ala83, and Lys85), and the hydrophobic pocket (Phe67 and Leu132). This broad interaction targets most of the enzyme active domains, except the activation loop. The comprehensive binding of 55E was used as a valuable template for designing novel inhibitors (Figure 1).

**Figure 1.**
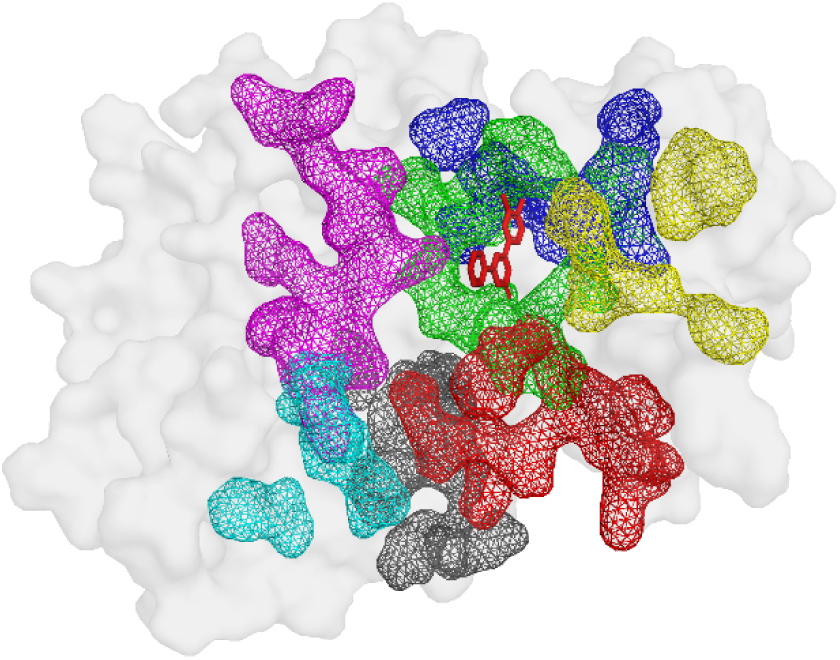
The surface representation of GSK-3β cavities including ATP binding site (green), substrate site (red), Axin/fratide site (cyan), as well as four allosteric sites (magenta, yellow, blue, and grey, respectively) in complex with 55E.

A total of 42 initial peptide (IPs) sequences were obtained from the kappa-casein sequence each consisting of 10 residues (Table 1). All peptides were docked against the binding site of 55E using the HADDOCK server. Given the larger size of the peptides relative to 55E, an expanded docking region was specified to accommodate the increased peptide length. The docking analysis identified IP8 (LRFFVAPFPE) as the top candidate for further improvement due to its most negative binding score, which suggests a highly favorable interaction with the enzyme’s active site (Figure 2).

**Table 1.**
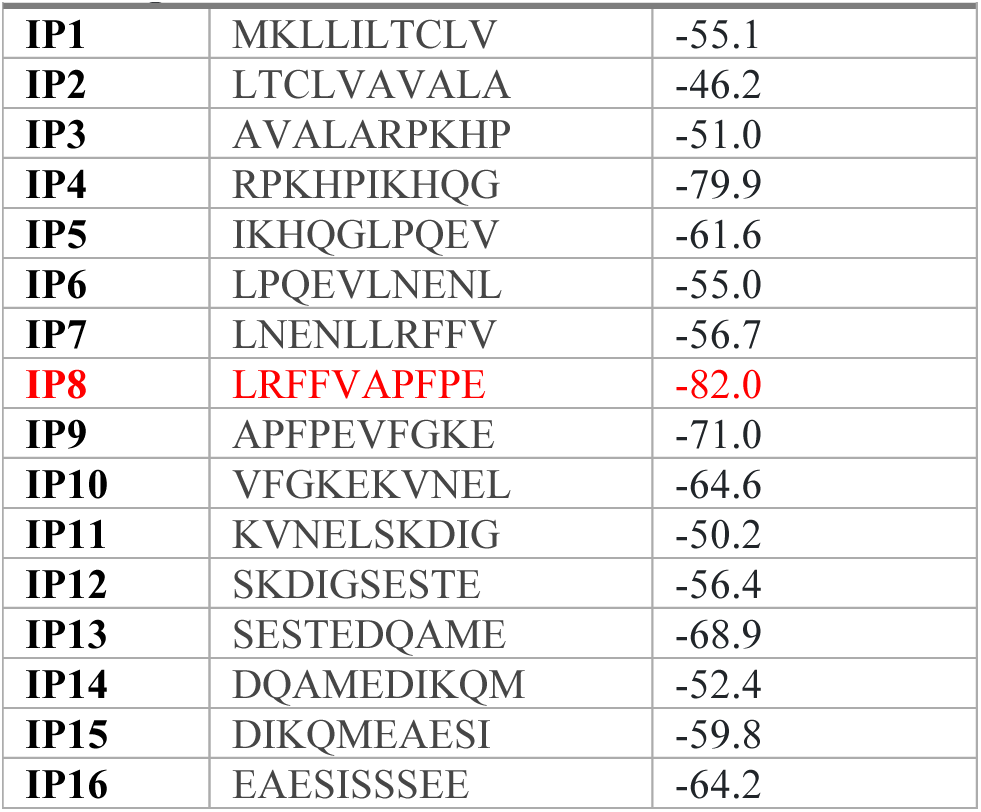
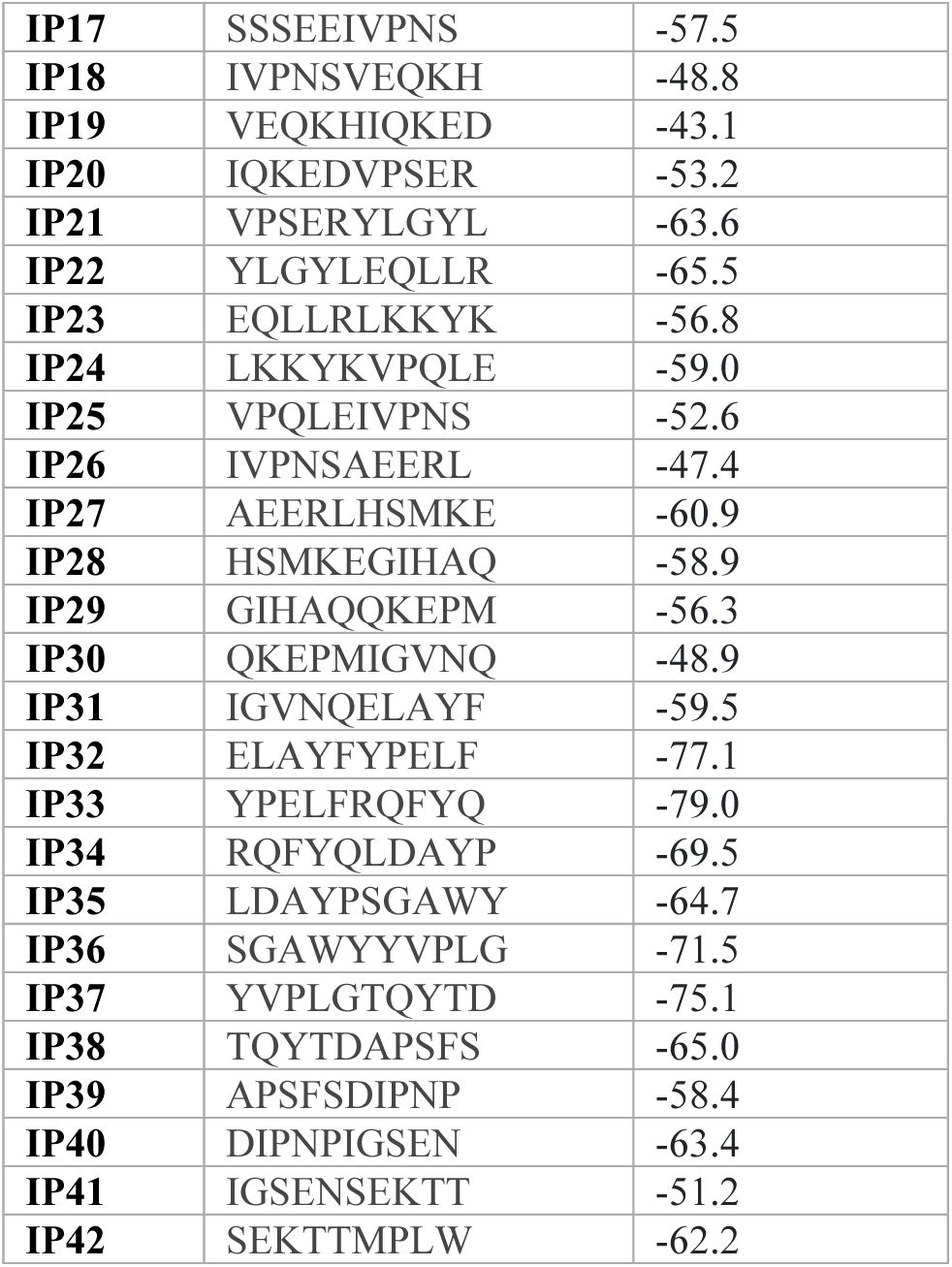
Template candidate sequences and their docking scores.

**Figure 2.**
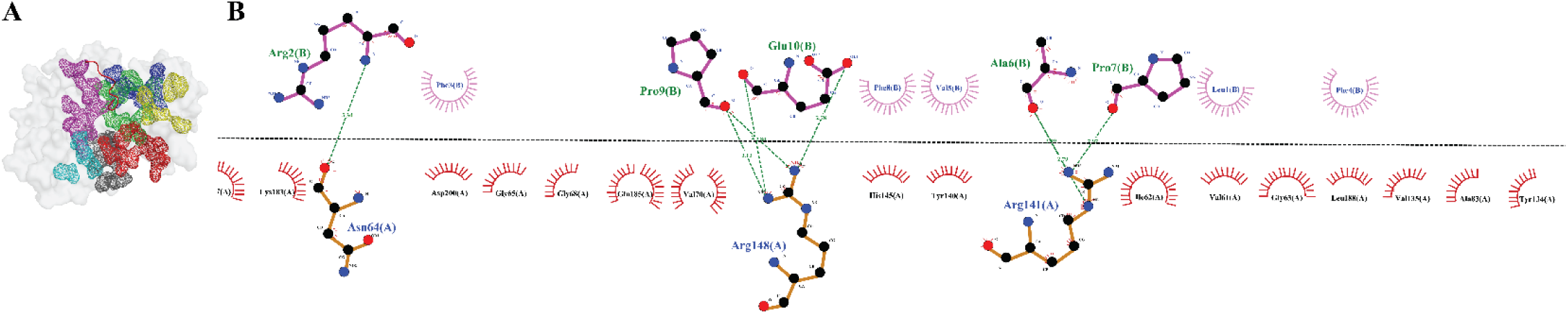
3D (A) and 2D (B) representation of the interaction between IP8 and the ATP binding pocket of GSK-3β, demonstrating the effective blockage of the binding site.

Although IP8 demonstrated strong binding, 150ns of MD simulation and the subsequent MMPBSA decomposition analysis revealed that the first three residues (LRF) had positive binding energies, suggesting that these residues contributed suboptimally to the binding free energy (Table 2).

**Table 2.** MMPBSA analysis of IP8 against GSK-3β.

| Table 2-MMPBSA analysis of IP8 against GSK-3β | |
| --- | --- |
| Residue No. | MMPBSA energy |
| 1 | 99.624 |
| 2 | 90.0431 |
| 3 | 0.437 |
| 4 | -4.9453 |
| 5 | -1.8042 |
| 6 | -1.3177 |
| 7 | -4.0347 |
| 8 | -6.1668 |
| 9 | -2.9492 |
| 10 | -235.294 |

Targeted modification of energetically unfavorable residues offers a promising strategy to convert unfavorable molecular interactions into more favorable ones, thereby improving the overall binding affinity. To enhance the binding affinity of IP8, we carried out a series of mutations targeting the first three residues. To do this, the MCSM (Membrane Curvature Server for Molecular Dynamics) tool was utilized to perform sequence mutation based on the Delta Delta G (ΔΔG) concept, which quantifies the change in binding free energy associated with amino acid substitutions. The results, as summarized in Table S1, identified several stabilizing mutations for each position, characterized by positive ΔΔG values which indicate a favorable impact on peptide binding stability. Subsequently, we generated all possible sequence combinations from the favorable mutated residues at each position to evaluate their inhibitory potential against the enzyme.

Following this, all forty mutant sequences were screened for allergenicity and toxicity, leading to the identification of fifteen mutated peptides (MPs) (MP4, MP5, MP6, MP11, MP12, MP15, MP17, MP18, MP19, MP20, MP27, MP31, MP32, MP37, and MP39) (Table S2).

To evaluate the inhibitory potential of peptide candidates, they were docked against the 5EE binding cavity. The docking results revealed that MP31, MP27, and MP39 exhibited the most favorable binding scores of -98.6, -96.5, and -91.5, respectively (Table 3), while the remaining peptides showed lower binding affinities compared to the template peptide (IP8).

**Table 3.** HADDOCK Docking results of Mutated Peptides and GSK-3β.

| Table 3-HADDOCK Docking results of Mutated Peptides and GSK-3β. | | | | | | |
| --- | --- | --- | --- | --- | --- | --- |
| Peptide ID | Docking Score | RMSD from the overall lowest-energy structure | Van der Waals Energy | Electrostatic Energy | Desolvation Energy | Buried Surface Area |
| **MP4** | -75.5 | 0.4 | -43.3 | -104.4 | 54.0 | 1219 |
| **MP5** | -67.4 | 2.5 | -42.0 | -61.1 | -19.0 | 1277.2 |
| **MP6** | -72.0 | 0.3 | -38.8 | -144.7 | -9.3 | 1291.3 |
| **MP7** | -74.5 | 1.3 | -43.7 | -110.5 | -16.6 | 1188.6 |
| **MP11** | -66.7 | 2.4 | -41.9 | -50.7 | -17.5 | 1092 |
| **MP12** | -68.5 | 2.3 | -44.3 | -44.0 | -19.7 | 1148.3 |
| **MP16** | -72.1 | 0.3 | -32.2 | -198.0 | -10.5 | 1230 |
| **MP17** | -71.2 | 0.2 | -43.1 | -69.0 | -18.8 | 1186 |
| **MP18** | -78.1 | 0.2 | -39.5 | -165.2 | -10.0 | 1297.1 |
| **MP19** | -73.5 | 0.2 | -34.0 | -194.1 | -7.9 | 1209.5 |
| **MP20** | -71.2 | 0.9 | -41.3 | -83.3 | -18.1 | 1200.7 |
| **MP27** | -96.5 | 0.3 | -49.6 | -131.6 | -24.0 | 1415.6 |
| **MP31** | -98.6 | 0.6 | -52.3 | -112.3 | -28.0 | 1436.4 |
| **MP32** | -79.0 | 0.3 | -41.5 | -145.3 | -18.0 | 1264.4 |
| **MP37** | -78.9 | 0.7 | -35.0 | -179.0 | -13.9 | 1267.5 |
| **MP39** | -91.5 | 0.6 | -50.0 | -106.5 | -23.2 | 1351.7 |

The top potent peptides have the following mutations:

1. MP27 (LRFFVAPFPE -> HLDFVAPFPE): L2H, R3L, F4D
2. MP31 (LRFFVAPFPE -> HPDFVAPFPE): L2H, R3P, F4D
3. MP39 (LRFFVAPFPE -> HADFVAPFPE): L2H, R3A, F4D

Notably, all three MPs share the F4D substitution, where Phe is replaced by Asp, resulting in the replacement of a hydrophobic residue with a negatively charged residue. Moreover, all final candidates share the substitution of Lys with His, indicating that a polar and less bulky residue with intermediate hydrophobicity at the N-terminal of the peptide may enhance binding affinity. Variations in the second position among the final MPs account for differences in their binding affinities.

The lower score of MP39 compared to the MP27 and MP31 indicates that a small residue in position 2 weakens binding activity, while a bulky residue like Lys can strengthen peptide-GSK-3β formation. Conversely, the lowest score of MP31 suggests that a moderate and rigid residue such as proline at the second position may significantly increase binding affinity.

Further examination of interactions between the MPs and GSK-3β shows that MP27 and MP39 form higher numbers of hydrogen bonds with the receptor, nine and eight, respectively, implying a potentially greater affinity towards the enzyme cavity (Figure 2A, C, D, and F). However, the results indicate that MP31, with the most favorable binding score, interacts with nineteen residues of the enzyme cavity, while MP27 and MP39 interact with sixteen residues (Figure 2 B and E). This suggests that a higher number of contact points collectively enhance binding affinity, even though they may contribute minimal binding energy individually. Moreover, a greater number of interacting residues facilitate more specific and unique interactions, aiding in discriminating the inhibitor from other potential ligands or substrates and increasing binding specificity. Complexes with higher contact points are expected to fit more precisely into the ATP binding cavity. Moreover, it is seen in Table 4 that MP27 and MP39 share similar interaction patterns but differ in residue-specific contacts, suggesting slightly different binding conformations. In contrast, MP31 shows unique interactions with Asp133 and Cys199, potentially providing a distinct binding mechanism compared to the others.

**Table 4.** The table of contacts between the peptides and GSK-3β residues. Hydrogen bonds are indicated with numbers in parentheses while the remaining residues indicate hydrophobic interactions.

| Table 4- The table of contacts between the peptides and GSK-3β residues. Hydrogen bonds are indicated with numbers in parentheses while the remaining residues indicate hydrophobic interactions. | |
| --- | --- |
| Peptide ID | GSK-3β residues |
| IP8 | Val61, Ile62, Gly63, Asn64(1), Gly65, Phe67, Gly68, Val70, Ala83, Val135, Tyr134, Tyr140, Arg141(2), His145, Arg148(4), Lys183, Gln185, Leu188, Asp200 |
| MP27 | Ile62, Gly63, Asn64(4), Gly65, Phe67, Val70, Tyr134, Val135, Thr138, Tyr140, Arg141(2), Arg144, His145, Arg148(4), Lys183(1), Gln185, Leu188 |
| MP31 | Ile62, Asn64(4), Gly65, Val70, Ala83, Asp133, Tyr134, Val135, Thr138, Tyr140, Arg141(2), Arg144, His145, Arg148(2), Lys183(1), Gln185, Leu188, Cys199 |
| MP39 | Ile62, Gly63, Asn64(2), Gly65, Val70, Tyr134, Val135, Thr138(1), Tyr140, Arg141(2), Arg144, His145, Arg148(2), Lys183(1), Gln185, Leu188 |

Following the study of the dynamic behavior of the enzyme and peptides over time, the RMSD analysis revealed significant fluctuations in GSK-3β, indicating that the protein experiences greater deviations from the initial conformation in the absence of a peptide ligand. These greater fluctuations suggest that the apo form of the peptide has more freedom to move. In contrast, when GSK-3β is bound to the peptides, less fluctuation is seen in RMSD plots, suggesting that peptide binding reduces the overall flexibility of the protein, probably leading to the restricted conformation. Notably, among the MPs, MP31 demonstrated the least fluctuation compared to other plots, suggesting that MP31 may limit the free movement of protein domains or side chains and make a more rigid structure more stable compared to the apo form. Additionally, Rg analysis, which measures the distribution of the atoms of a molecule around its center of mass, showed that the enzyme in its apo- and MP39-bound states exhibited more extended conformations compared to MP31 and MP27 (Figure 3B). The constant Rg throughout the simulation period indicates the absence of any unfolding events.

**Figure 3.**
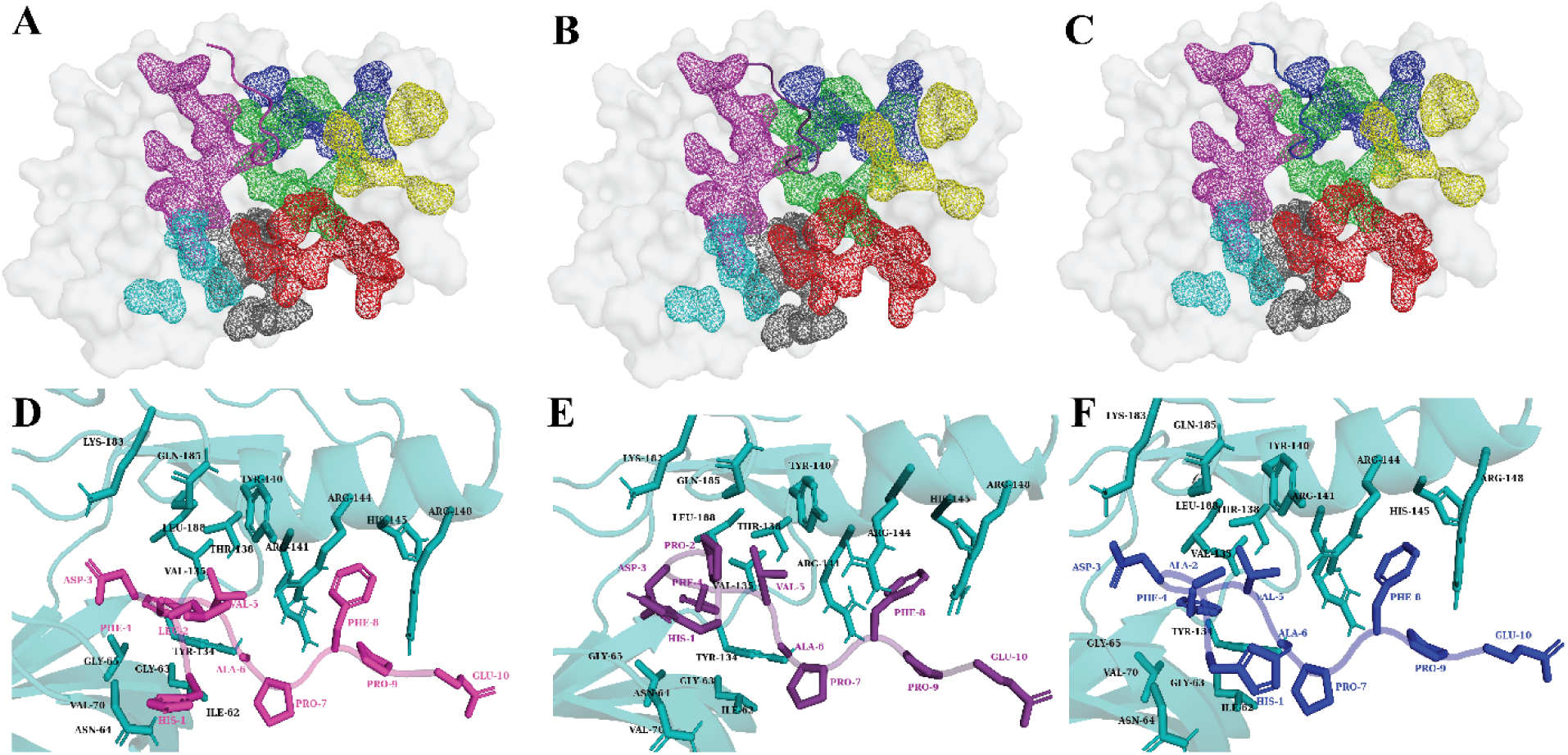
The 3D representation of the interacted residues of GSK-3β with the peptide inhibitors MP27(A), MP31(B), and MP39(C), visualized using PyMOL. Interacting residues of GSK-3β, MP27, MP31, and MP39 are shown in light blue, magenta, deep violet, and deep blue, respectively. 2D diagram of interactions designed peptide inhibitors, MP27(D), MP31(E), and MP39(F), with the GSK-3β enzyme (chainA). The diagrams highlight key hydrogen bonds, hydrophobic interactions, and other significant contacts between the peptides and the active site residues of GSK-3β.

The SASA values for the peptide-bound complexes are consistently lower than those of the apo form, with an average of 183.17, 180.2, 182.3, and 181.68 nm² for apo-protein, MP27, 31, and 39 respectively. This suggests that peptide binding induces a more compact conformation, reducing the solvent exposure of the protein. The reduced SASA in the peptide-bound systems indicates that the peptides promote a tighter packing of the protein structure. This compaction may shield hydrophobic residues from solvent exposure and restrict the flexibility of critical loops or regions near the active site, potentially obstructing substrate access or product release, which are crucial for enzyme activity. In the apo form (light blue), the protein is more solvent-exposed, facilitating substrate interaction and catalytic efficiency (Figure 3C).

This finding aligns with the backbone RMSF analysis, which measures the average fluctuations of individual backbone atoms within each residue. As depicted in Figure 3D, most residues of GSK-3β exhibited similar backbone volatility in the peptide-bound state compared to the apo-enzyme, with notable exceptions observed in Phe67, Gln365, and Asn370, which display higher flexibility in the apo state (Figure 4D).

**Figure 4.**
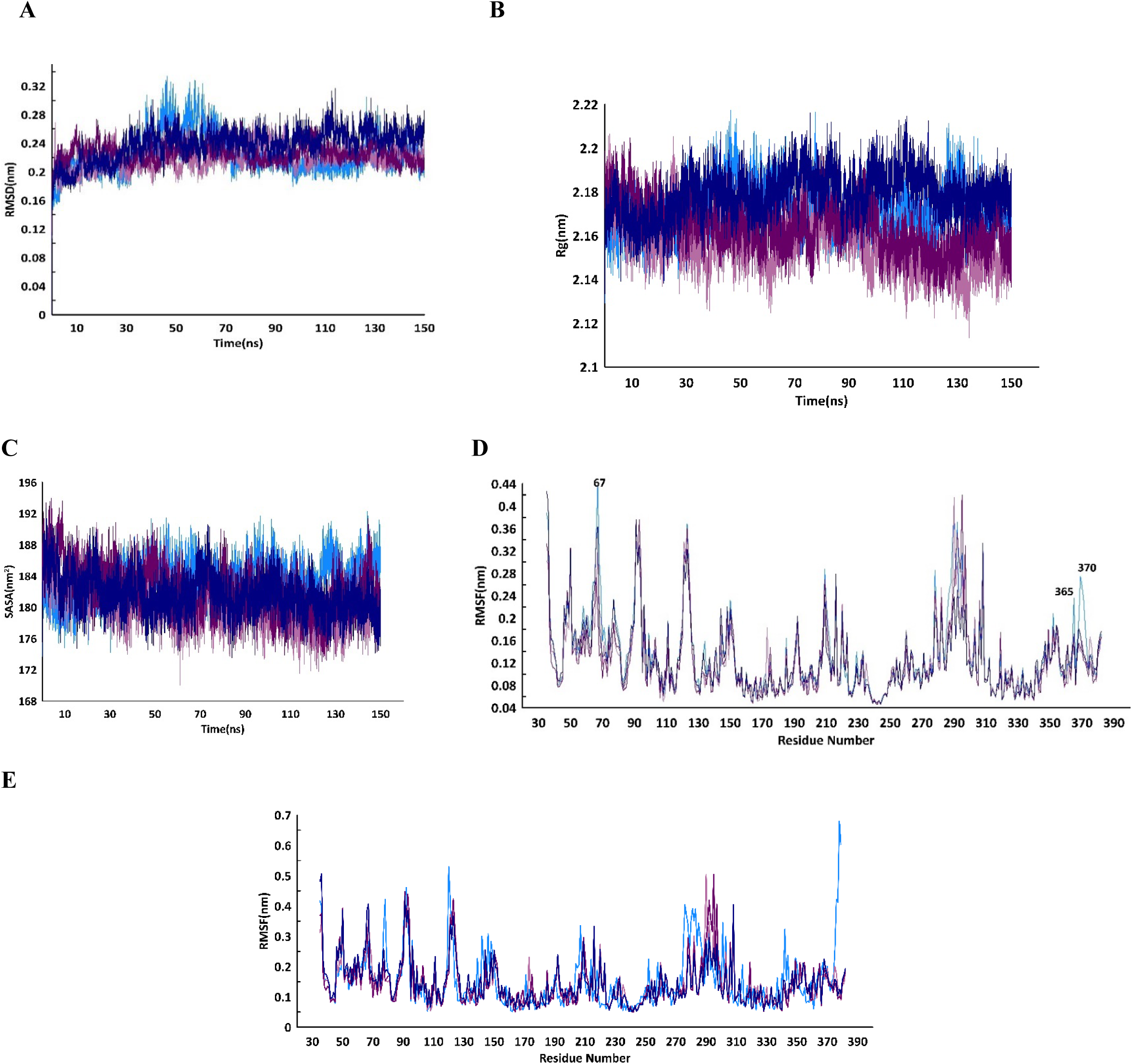
Comparative dynamic analysis of GSK3-beta in its apo (light blue) form and in the MP27-(magenta), MP31-(deep violet), and MP39-(deep blue) bound complexes. (A) RMSD, (B) Rg, (C) SASA, (D) backbone RMSF, and (E) side chain RMSF.

When comparing this to the side-chain RMSF analysis (Figure 4E), it is evident that the backbone fluctuations are significantly lower than those of the side chains. This is expected, as the backbone provides a structural framework of the protein, while side chains are inherently more dynamic. Notably, the side chains in the peptide-bound enzyme exhibit reduced flexibility, indicating a higher degree of dynamic behavior of the apo-enzyme. This observation provides further evidence that peptide binding restricts the flexibility of GSK-3β, both in the backbone and side chains.

Furthermore, the analysis reveals that the N-terminal domain of the protein (residues1-134) had lower fluctuations compared to the C-terminal domain (residues 135-382), highlighting the differential dynamic behavior across the two regions. The binding of peptides further reduces these fluctuations in both domains, reinforcing the idea that peptide binding stabilizes the overall structure and limits the intrinsic movements of both the backbone and side chains.

The behavior of hydrogen bonds was observed to be highly dynamic over the simulation period. Despite the lower count of hydrogen bonds, MP31 showed greater potency against the enzyme compared to MP27 and MP39, which exhibited higher hydrogen bond counts. This discrepancy can be attributed to the stability of the H-bonds between the peptide inhibitors and the GSK-3β ATP binding cavity. The MD results showed that the hydrogen bonds in MP27 and MP39 were predominantly unstable, while MP31 exhibited more stable H-bonds (Figure 5).

**Figure 5.**
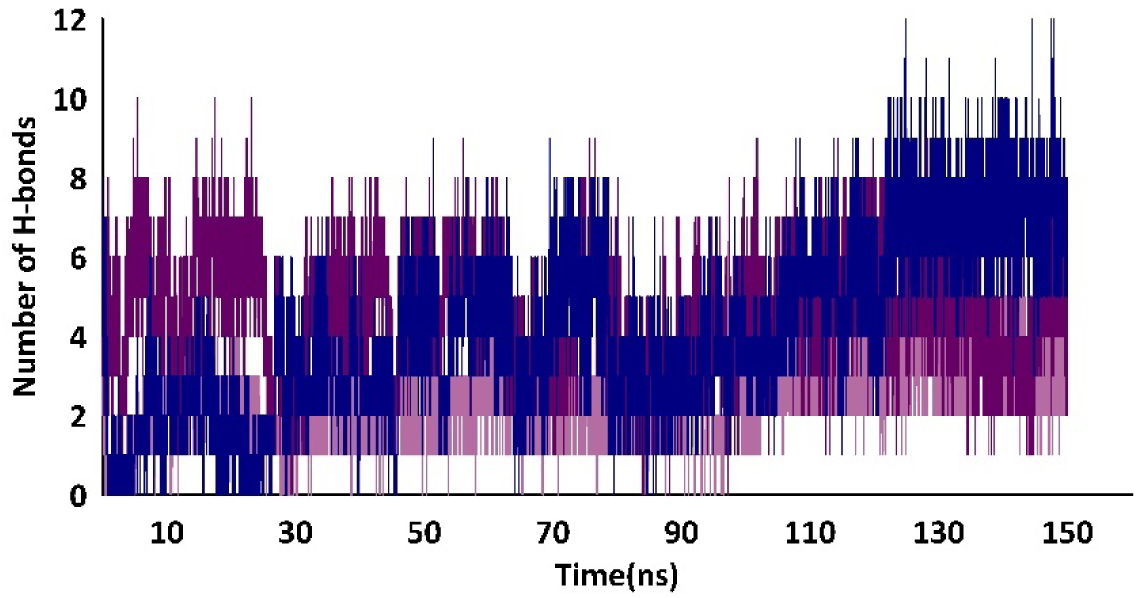
Hydrogen bond analysis of GSK3-beta in its apo (light blue) form and the MP27-(magenta), MP31-(deep violet), and MP39-(deep blue) bound complexes.

To analyze the movements and structural changes in the GSK-3β structure during the simulations in both unbound (apo) and bound states, PCA was conducted on the final 10 ns of each trajectory. The GSK-3β enzyme displayed two distinct and widely dispersed clusters, reflecting the diverse conformations assumed by the enzyme over time. This dispersion indicates a high level of flexibility due to the broad distribution (Figure 6A). Conversely, upon binding of MPs, the protein exhibited multiple clusters, illustrating various conformations adopted by the protein in its complex states. Analysis of the temporal progression revealed that all clusters occurred at least once during the final 10ns of the simulations. Moreover, the narrower distribution of MPs-bound complexes indicates limited conformational changes, suggesting a more constrained conformational space (Figures 6B, C, and D). These findings demonstrate that the presence of MPs stabilizes multiple conformations in the protein structures. Moreover, it was observed that the peptide-bound enzyme exhibited a reduced range of motion along both Principal Component 1 (PC1) and Principal Component 2 (PC2) compared to the apo form. This reduction in conformational flexibility suggests that the binding of peptides imposes structural constraints on the enzyme, limiting its ability to explore a diverse range of conformations. In the apo form, the enzyme’s broader range of motion along PC1 and PC2 typically reflects the dynamic behavior necessary for efficient catalytic activity, including substrate binding, product release, and allosteric regulation. However, peptide binding appears to restrict these essential conformational fluctuations, leading to a more rigid and potentially less functional enzyme state.

**Figure 6.**
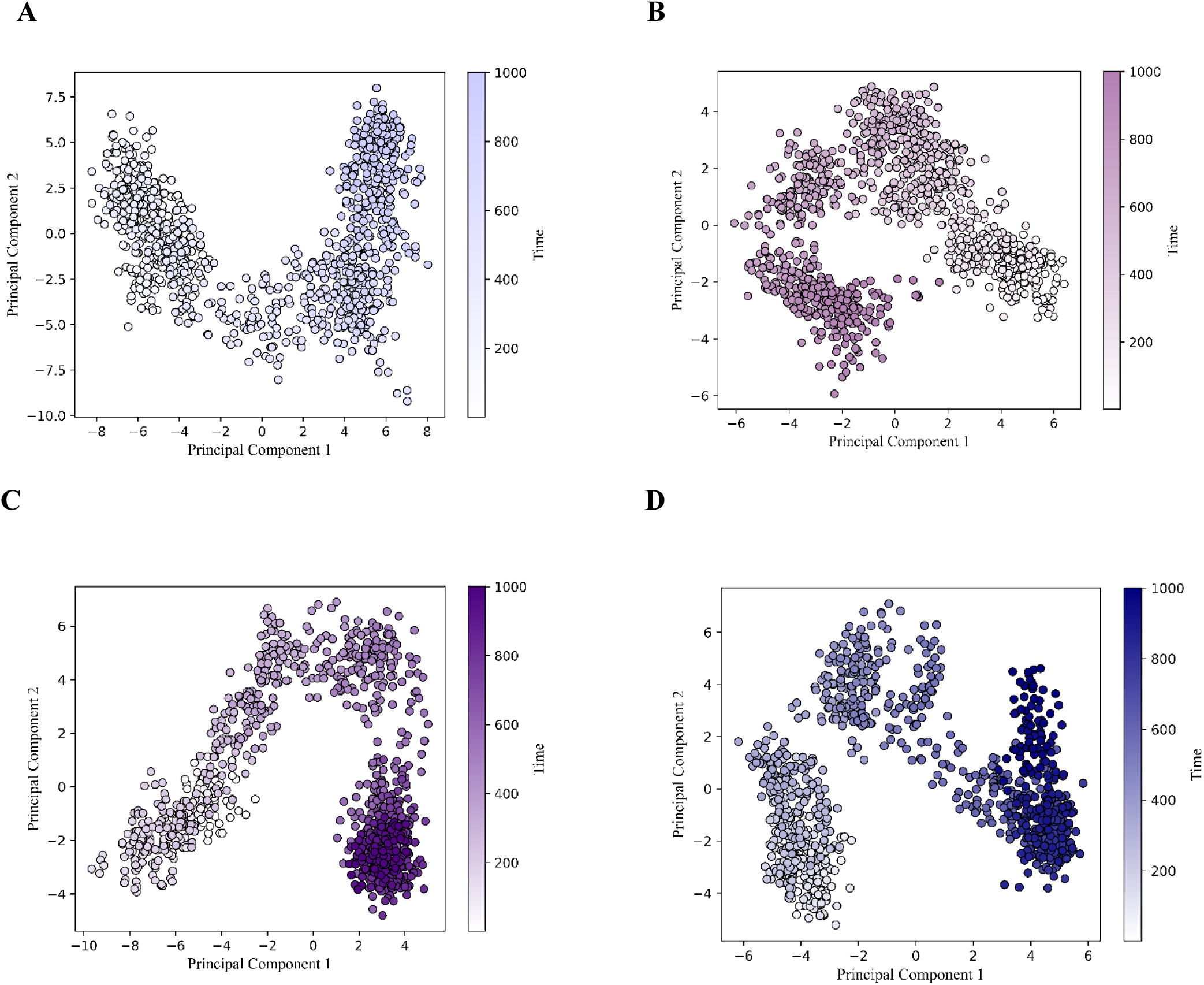
Principal Component Analysis (PCA) of Protein Dynamics PCA visualization depicting the conformational dynamics of the protein in Apo (A) as well as MP27- (B), MP31- (C), and Mp39- (D) states derived from the last 10 ns (1000 frames) of the trajectories.

The structural differences between PCA clusters in the simulations were quantified by calculating the pairwise RMSD values between the centroids of each cluster. As shown in Table S3, the RMSD values for GSK-3β in its apo form are notably larger (0.230 nm between clusters A and B), reflecting greater conformational flexibility. In contrast, peptide-bound systems (GSK-3β-MP27, GSK-3β-MP31, and GSK-3β-MP39) exhibit smaller RMSD values between clusters, with pairwise differences ranging from 0.141 nm to 0.175 nm. These results indicate that the peptides induce structural rigidity in GSK-3β, reducing the magnitude of conformational changes compared to the apo state. Thus, the differences in RMSD values suggest a clear trend of reduced conformational variability upon peptide binding, supporting the hypothesis that peptide interaction induce rigidity within the protein structure.

The data presented in Table 6 illustrates the mean MMPBSA binding energies for three final candidate peptides in complex with the enzyme GSK-3β. The binding energies are categorized into van der Waals energy, electrostatic energy, polar solvation energy, SASA energy, and total binding energy. Notably, the van der Waals interactions are most pronounced for MP39, indicating its superior hydrophobic interactions with GSK-3β compared to the other peptides. MP39 exhibits the highest electrostatic energy contribution, implying its formation of the most robust electrostatic interactions with the enzyme, followed by MP31 and then MP27. Moreover, the polar solvation energy is most elevated for MP39, signifying a greater desolvation penalty, which is characteristic of more polar or charged interactions. The SASA energy is most favorable for MP39, suggesting it buries a larger surface area upon binding in comparison to the other peptides. When considering the total binding energy, which encompasses all interaction components, MP31 emerges as having the most favorable overall binding energy, followed by MP27. Despite MP39 displaying strong individual interactions, its total binding energy is less favorable due to the high polar solvation energy. In conclusion, MP31 demonstrates the most favorable total binding energy with GSK-3β, indicating it is the strongest binder overall. MP39, while exhibiting strong van der Waals and electrostatic interactions, incurs a high desolvation penalty, resulting in a less favorable total binding energy. MP27, although showing moderate interaction energies, still exhibits strong binding affinity, albeit not as high as MP31.

**Table 6.** Average MMPBSA Binding Energies of MP27, MP31, and MP39 with GSK-3β (kJ/mol).

| Table 6- Average MMPBSA Binding Energies of MP27, MP31, and MP39 with GSK-3β (kJ/mol). | | | |
| --- | --- | --- | --- |
| Energy | MP27 | MP31 | MP39 |
| Van der Waal energy | -202.435 | -215.190 | -218.363 |
| Electrostatic energy | -149.886 | -376.339 | -680.489 |
| Polar solvation energy | 229.568 | 447.732 | 824.475 |
| SASA energy | -24.693 | -28.901 | -34.616 |
| Total Binding energy | -147.446 | -172.698 | -108.993 |

The utilization of inhibitors targeting the ATP-binding domain of multifunctional enzymes such as GSK-3β presents various challenges, notably the risk of adverse effects resulting from complete enzyme inhibition. However, Tideglusib serves as a notable instance of an ATP-binding pocket inhibitor that has progressed to clinical trials (31–33). Additionally, the utilization of peptide-based medications in addressing central nervous system (CNS) disorders needs further caution due to the limited permeability of the peptide drugs across the blood-brain barrier (BBB).

## Conclusion

This study demonstrates the potential of kappa casein-derived peptides, particularly MP31 (HPDFVAPFPE), as effective inhibitors of GSK-3β, a critical target in Alzheimer’s disease (AD) progression. The comprehensive analysis, including docking, molecular dynamics (MD) simulations, and structural stability assessments, highlighted MP31’s superior binding affinity, stability, and specificity for GSK-3β. PCA further confirmed that MP31 induced fewer conformational changes in the enzyme, emphasizing its role in stabilizing multiple protein conformations. Additionally, MP31’s favorable total binding energy, supported by strong van der Waals and electrostatic interactions, positions it as the most potent inhibitor of the candidate peptides. These findings support the development of MP31 and related peptides as therapeutic candidates for AD by inhibiting GSK-3β and potentially mitigating key pathological processes. Future studies will focus on in vitro and in vivo validation to further assess the therapeutic potential of these peptides in AD treatment.

